# STAT3 regulates cytokine production downstream of TNFR1 in part by inducing expression of TNFAIP3/A20

**DOI:** 10.1101/2022.01.19.476699

**Authors:** Ricardo Antonia, Eveliina Karelehto, Kan Toriguchi, Mary Matli, Robert Warren, Lawrence M. Pfeffer, David B. Donner

## Abstract

**Background:** Previous work demonstrated that the Signal Transducer and Activator of Transcription 3 (STAT3) is activated downstream of the Type 1 TNF Receptor. However, whether and how STAT3 regulates gene expression downstream of TNFR1 has not been elucidated.

**Methods:** Global transcriptome analysis by RNA sequencing was performed in wild type and STAT3 knockout mouse embryonic fibroblasts (MEFs) stimulated with TNF. The fold changes in gene expression were assessed bioinformatically. Results of the RNA sequencing were validated at the protein level by using multiplex cytokine assays and immunoblotting.

**Results:** Stimulation of MEFs with TNF or an agonist antibody to TNFR1 activated STAT3, and this was inhibited by pharmacological inhibition of Jak2 and cSrc. At 4 hours after TNF stimulation, STAT3 knockout MEFs had a greater level than WT MEFs of induction of the chemokines Ccl2, Cxcl1 and Cxcl10 at the RNA and protein levels. Mechanistically, this was due STAT3 promoting the expression of Tnfaip3/A20, a ubiquitin modifying enzyme that inhibits inflammation, in wild-type MEFs at early timepoints after TNFR1 stimulation. In STAT3 knockout MEFs TNF failed to induce the expression of Tnfaip3/A20 or GM-CSF when acting through TNFR1. Expression of A20 into STAT3 knockout MEFs suppressed cytokine expression.

**Conclusion:** STAT3 limits the induction of Ccl2, Cxcl1 and Cxcl10 in response to TNFR1 activation by promoting the expression of Tnfaip3/A20. On the other hand, STAT3 promotes the expression of GM-CSF in response to TNFR1 stimulation. These results show that STAT3 modulates inflammatory signaling by TNF in normal cells.

## Introduction

TNF plays a central role in integrating and amplifying the host response to infection and malignancy. Through interactions with macrophages, fibroblasts and endothelial cells, TNF promotes and coordinates the immune response, local inflammatory processes, and wound repair as it stimulates the synthesis of various other cytokines, such as interleukin 1, interleukin 6, and granulocyte-macrophage colony-stimulating factor (1–4).

Cellular responses to TNF are initiated by two receptors, the type 1 TNFR (TNFR1) and the type 2 TNFR (TNFR2)(*1–3*) Most TNF actions are mediated by TNFR1, which contains a death domain that fosters protein-protein interactions (*1–3*). Although TNFR1 does not have intrinsic tyrosine kinase activity, various phosphorylation reactions are induced by TNF and necessary for its biologic effects(*4–8*). These phosphorylation events are associated with alterations of cellular sensitivity to TNF-mediated cytotoxicity (*7, 9*) Inhibitors of protein tyrosine kinases suppress TNF-stimulated DNA fragmentation(*10*), NF-κB activation, and the expression of endothelial cell adhesion molecules(*11, 12*). The priming of neutrophils by TNF is also accompanied by tyrosine phosphorylation events that direct the cells to undergo a respiratory burst (*4–6*) Our laboratory has shown that TNF induces the tyrosine phosphorylation of a group of cytoplasmic proteins, including PI3K and insulin receptor substrate-1,(*13*) promotes NF-κB activation (*14–16*) and modulates the responsiveness of cells to insulin.(*17*)

Cytokine receptors that do not possess tyrosine kinase activity use nonreceptor tyrosine kinases, such as members of the Jak and Src family, with which they associate to initiate signaling.(*18*) We and others previously showed that Jak2 and cSrc associate with TNFR1 and that through these tyrosine kinases TNFR1 induces the activation of STAT proteins, including STAT3. (*19–21*) However, the role of STAT3 downstream of TNFR1 in promoting any of the various functions of TNF, particularly those associated with inflammatory processes, is much less characterized.

STAT3 is a transcription factor that is activated by cytokine and growth factor receptors, and has many important roles in normal physiology and disease (*22*). Insofar as TNFR1 signaling is concerned we previously showed that in some cell types this is through the formation of TNFR1 complexes with Jak2 which mediates the phosphorylation of tyrosine-705 on STAT3.(*19*) Additionally, many potent STAT3 agonists (notably IL-6, LIF) are downstream transcriptional targets of TNFR1 and induction of these genes further amplifies the levels of STAT3 activation downstream of TNFR1. (*23*)

While the roles of other signaling molecules downstream of TNFR1, such as NF-κB (*24, 25*) and Jun (*26*), are well characterized, the role of STAT3 is less well characterized, although STAT3 promotes mitochondrial respiration in TNF-treated mouse embryonic fibroblasts (MEFs) (*21*). In addition, the role of STAT3 in response to TNFR1 stimulation in regulation of gene expression has not been characterized at a global level. Depending on the stimulus and the cellular context STAT3 either promotes inflammation or inhibit pro-inflammatory gene expression. What role STAT3 plays in inflammatory gene expression downstream of TNFR1 is unclear as STAT3 activation induced by Interleukin-6 (IL-6) promotes inflammation, while activation induced by Interleukin-10 inhibits inflammation (*27, 28*).

To shed light on the role of STAT3 in gene expression after TNFR1 stimulation, we analyzed the changes in the transcriptome that occurred in response to TNFR1 stimulation in non-transformed cells, mouse embryo fibroblasts.

## Results

### TNFR1 activates STAT3 in MEFs through Jak/Src

Our previous studies showed that TNF induces STAT3 activation in various transformed cell lines through activation of TNFR1-associated Jak2. The present study expands on these observations by testing whether and how TNF would induce activation of STAT3 in MEFs and then testing for the significance of this signaling event.

Depending on the cell type used, c-Src or Jak2 activate different STAT proteins(*29, 30*). We tested whether activation of TNFR1 induced STAT3 activation in MEFs. Time-course experiments showed that low concentrations of TNF*α* (0.1nM) induce a biphasic induction of STAT3 phosphorylation (Fig. 1A). One burst of STAT3 phosphorylation occurred at 5-15 min after TNF*α* treatment, and another after 45 min. This biphasic signaling is similar to what is observed for TNF-induced NF-κB signaling.(*31*) We confirmed that STAT3 induction was mediated through TNFR1 as a TNFR1 agonistic antibody phenocopied STAT3 phosphorylation and activation by TNFα (Fig. 1A). In MEFs STAT3 activation was dependent on both Jak and Src, because Jak or Src inhibition diminished TNFα induced STAT3 phosphorylation (Fig. 1B).

**Figure 1:**
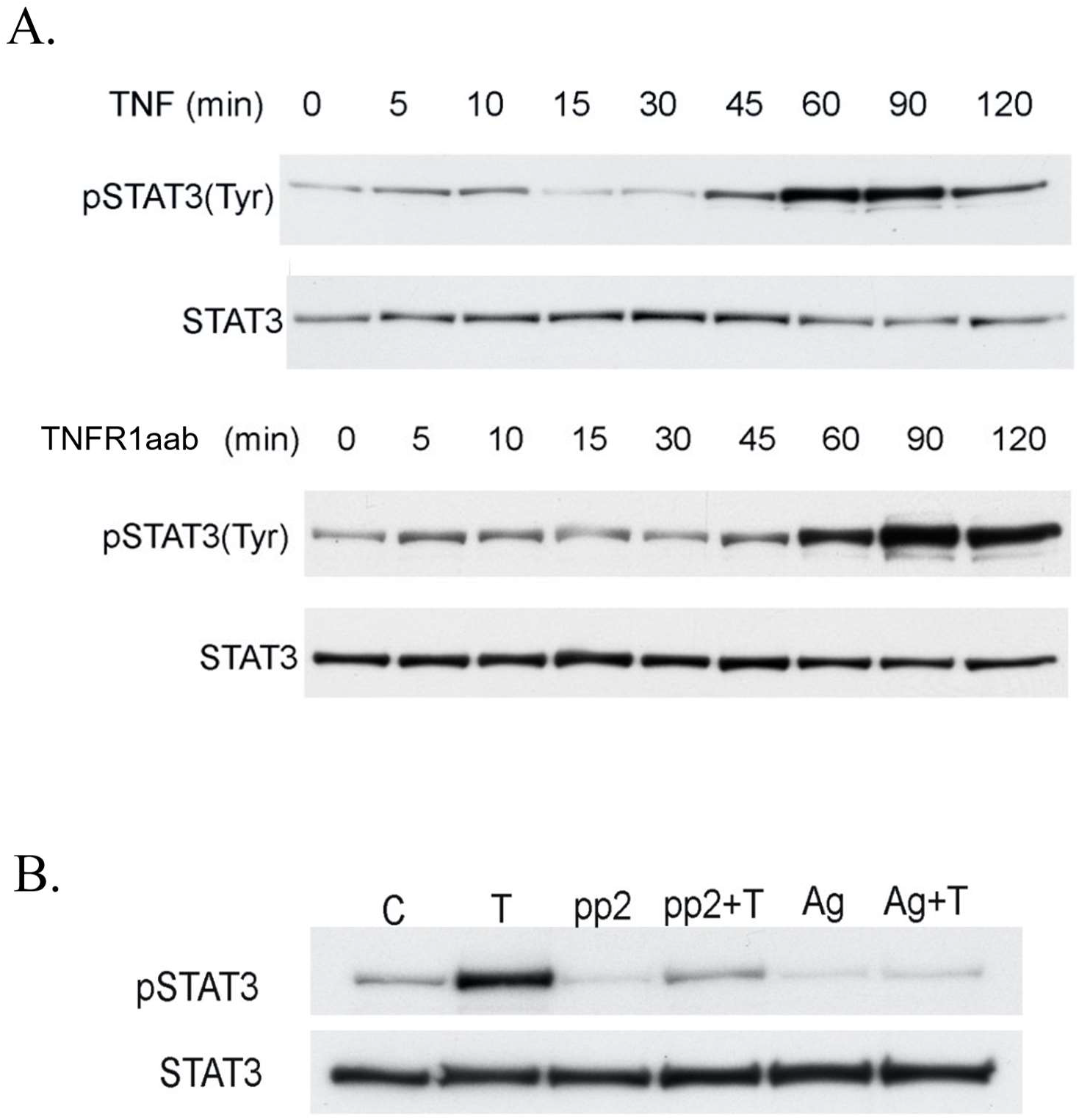
STAT3 is activated by TNFR1 in a Jak/Src dependent manner A. MEFs were stimulated with TNF-α (top) or a TNFR1 agonist antibody (bottom) for the indicated times before being lysed and analyzed by immunobloting with either a phospho-(Y705) or total STAT3 antibody. B. MEFs were pre-treated for 1 hr with DMSO, AG490 (50 uM), or PP2 (10 uM), then stimulated with TNF for 60 min, before immunobloting with phospho(Y705) or total STAT3 antibody.

### Characterization of transcriptomic changes in Wt and Stat3−/− MEFs 4 hrs after TNFα treatment

To characterize the functional role of STAT3 activation induced by TNF, we performed RNA sequencing on RNA prepared from wild type (*Wt*) and STAT3 knockout (*Stat3−/−*) MEFs that were either unstimulated or stimulated with TNFα for 4 hours. After mapping and quality control steps, the fold change in gene expression was assessed for the response to TNFα relative to the unstimulated control for each MEF genotype (Fig. 2A). The resulting volcano plots of gene expression changes showed that TNF induced a greater fold change in gene expression in *Stat3−/−* MEFs as compared to Wt MEFs.

**Figure 2:**
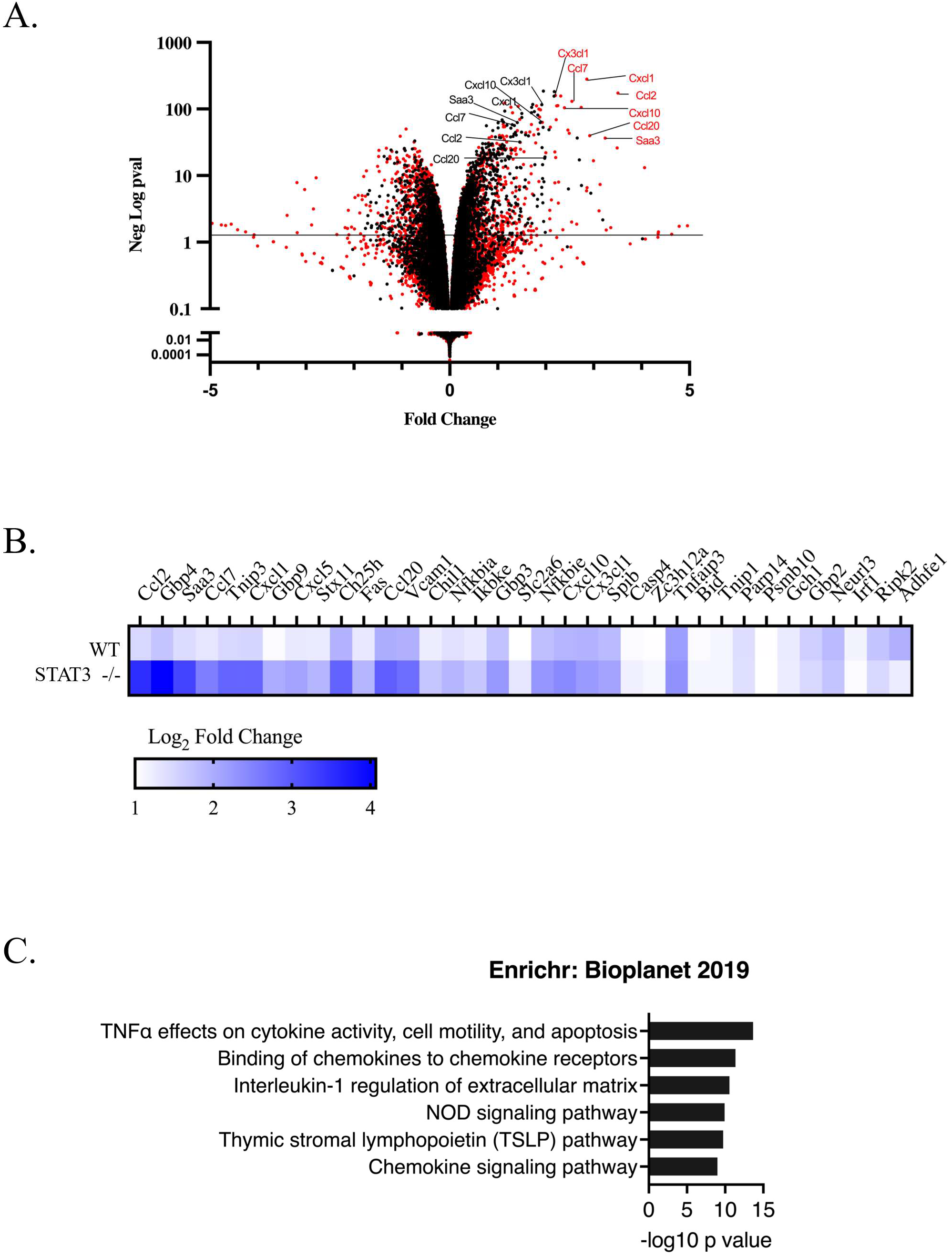
Increased chemokine/cytokine expression in STAT3−/− MEFS in response to TNFR1 stimulation with TNFα for 4 hours A. Volcano plot showing the differentially expressed genes in WT and matched STAT3−/− MEFs stimulated with TNFα for 4 hours B. Heat map of the validated TNFα targets in WT or STAT3−/− MEFs C. ENRICHR pathway analysis of STAT3 repressed genes

To further characterize TNF-induced genes, we next sought to identify a group of validated TNFα target genes to minimize potential false positives. We therefore used RNA sequencing data from Wt MEFs derived from different genetic backgrounds that were stimulated with TNFα or with a TNFR1 agonist antibody (TNFR1aab). The overlapping group of genes that were induced in each MEF genotype by TNF or the TNFR1aab were included in the final “validated” list of TNFα target genes.

Of the 35 genes that were validated as TNFR1 dependent genes, the majority (23 genes) had a greater induction in the *Stat3−/−* MEFs (Stat3 repressed genes). In addition, 2 genes had lower induction in the *Stat3−/−* MEFs (*Stat3* induced genes), while 10 genes had similar levels of induction (*Stat3* independent genes) in response to TNFα (Figure 2B). Since most of the validated genes were repressed by *Stat3*, we focused on this group of genes. Enrichr (*32–34*) analysis of the Stat3 repressed genes indicated that these were characteristic of the “TNFα Signaling via NF-κB” signature, and the “Binding of chemokines to chemokine receptors” signature (Figure 2C) (32–34). Overall, transcriptomic analysis indicated that Stat3 down-regulates the expression of many but not all TNFα regulated genes.

### Stat3 represses the secretion of CCL2, CXCL1, and CXCL10 and promotes secretion of GM-CSF downstream of TNFR1

We next determined if the observations made at the gene expression level resulted in changes of protein abundance and secretion. Since TNFα can bind to TNFR1 or TNFR2, we also validated that the observations were specific to TNFR1 by comparing TNFα stimulation with TNFR1aab stimulation. Since the top matching signature for the *Stat3−/−* repressed genes in response to TNFα was the “Binding of chemokines to chemokine receptors” and many of the genes in this group encoded secreted cytokines, we performed multiplex assays using arrays composed of a panel of 44 murine cytokines. For this assay, cell culture supernatants of *Wt* and *Stat3−/−* MEFs were collected 4 hr after mock or TNFα stimulation, or after treatment with the TNFR1aab. The supernatants were then analyzed using the multiplex cytokine assay.

To avoid false positives, we set a cut-off of at least a two-fold change for a result to be considered a true positive. Consistent with the RNA sequencing analysis, Ccl2/Mcp1, Cxcl1/KC, and Cxcl10/IP10 were induced to a greater degree in the *Stat3−/−* versus *Wt* MEFs in response to TNFα or the TNFR1aab (Fig. 3A). Thus, RNA sequencing and assays of cytokine protein expression were concordant in that both showed that *Stat3−/−* MEFs had greater levels of cytokine gene expression and secretion downstream of TNFR1 as compared to their Wt counterparts. These observations show that STAT3 plays a role in limiting the secretion of specific cytokines downstream of TNFR1.

**Figure 3:**
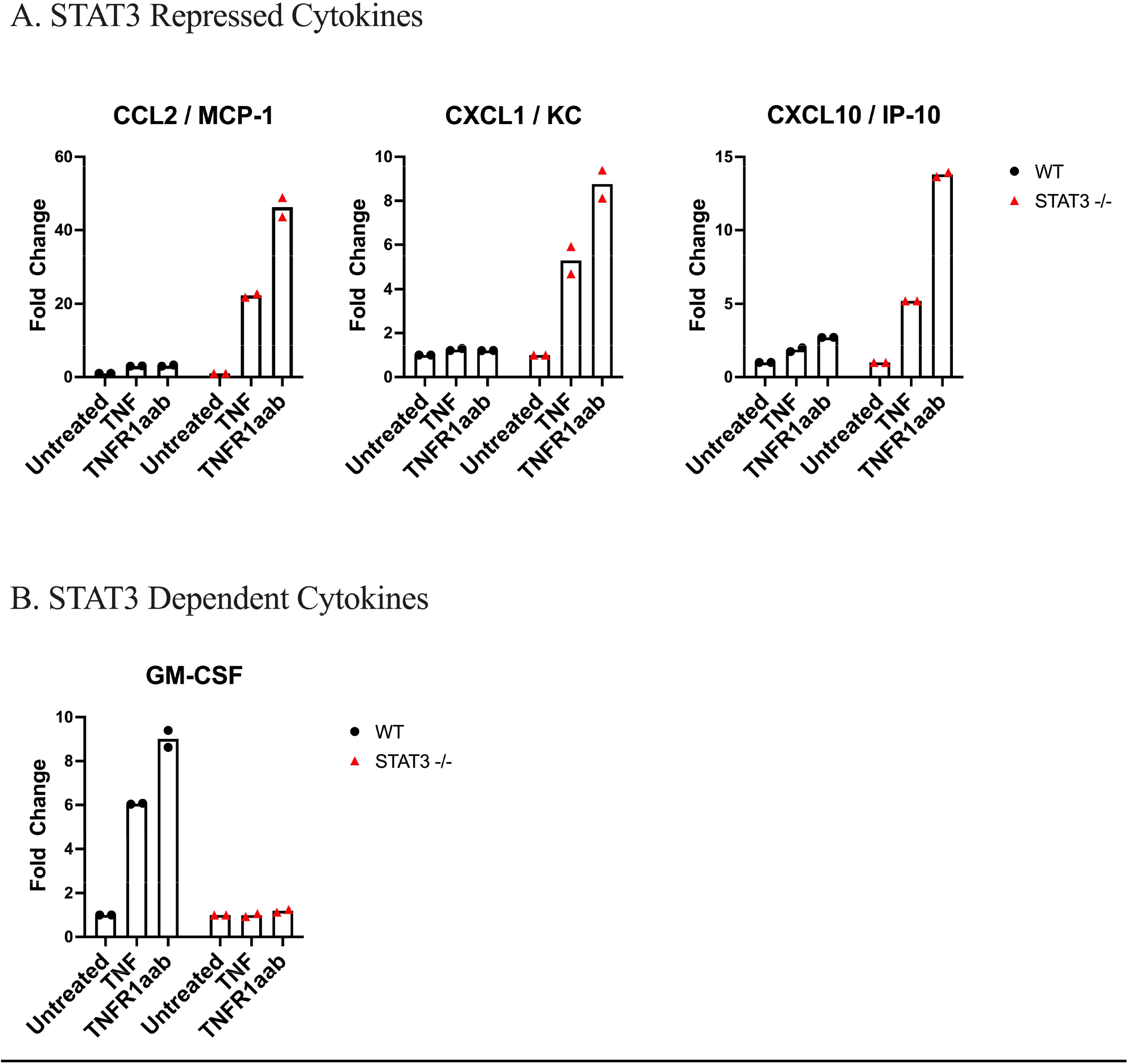
Protein level validation of the above RNA sequencing using multiplex cytokine assays. Wt or STAT−/− MEFs were treated with vehicle, TNF or TNFR1aab for 4 hours at which time supernatants in the cell cultures were collected. Cytokine expression in the supernatants was assayed using the multiplex cytokine array as described in the methods. Panel A. Cytokines whose expression was greater in Stat3−/− MEFS (i.e. suppressed by STAT3). Panel B. Cytokine whose expression was diminished in Stat3−/−MEFs (i.e. induced by STAT3).

Of the cytokines in the array panel, GM-CSF was the only cytokine or growth factor that was induced to a greater extent in *Wt* than in *Stat3−/−* MEFs, indicating that its induction was *Stat3* dependent. Since GM-CSF was not detected by RNA sequencing in unstimulated cells, a fold-change could not accurately be calculated. Nonetheless, protein-based assays show that GM-CSF secretion is STAT3 dependent in response to TNF-α (Fig. 3B).

The remaining cytokines in the panel, such as LIF, were either induced to a similar degree in *Wt* and *STAT3−/−* MEFs or had inconsistent patterns upon stimulation with TNF and the TNFR1aab, indicating that they were either not regulated by STAT3 or not regulated by TNFR1.

### STAT3 activation induces the expression of A20/Tnfaip3 thereby suppressing Ccl2, Cxcl1, and Cxcl10 expression

We next sought to understand the mechanism whereby STAT3 could repress the expression of Ccl2, Cxcl1, and Cxcl10. The experiments shown in Fig. 3 were performed after 4 hr of TNF stimulation. We hypothesized that STAT3 may induce a negative regulator of TNFR1 at a relatively early timepoint, and that failure to induce this regulator could explain the excess cytokine production in the *Stat3−/−* MEFs.

To identify genes regulated at an early timepoint by activated STAT3, we performed RNA sequencing of the *Wt* and *Stat3−/−* MEFs that were either unstimulated or stimulated for 30 min with TNFα, and the fold change was calculated for each gene and expressed in a volcano plot (Figure 4A). In contrast to the gene expression changes at 4 hr, the magnitude of the changes induced by TNFα at 30 min were similar in the *Wt* and *Stat3−/−* MEFs. However, genes that were repressed in the *Stat3−/−* as compared to *Wt* MEFS by TNFα (Figure 3B), included *Iex-1*, *Zfp36/TTP*, and *A20/Tnfaip3* which are established negative regulators of the TNFR1 pathway (Figure 4B)(*35–37*). The only other negative TNFR regulator that was up regulated was Nfkbia (IκBα), but this did not appear to be regulated by Stat3. To validate the role of Stat3 activation observed in our RNAseq data, we performed western blots on *Wt* MEFs stimulated with the TNFR1aab, in the presence or absence of the STAT3 inhibitor Stattic (*38*). Western blots failed to detect Iex-1 and Zfp36 (data not shown), and we suspect that this was due to the low concentration of TNFR1aab used in these experiments or that the protein abundance of these genes is regulated post-transcriptionally. Since these proteins were not detected, we reasoned that they would be unlikely to be responsible for the increased cytokine production observed in the *Stat3−/−* MEFs. In contrast, A20/Tnfaip3 was induced by the TNFRaab and this induction was blocked by Stattic (Figure 4C). A20/Tnfaip3 is a deubiquitinating enzyme that inhibits TNFR1 signal transduction (*37*). We therefore hypothesized that STAT3 was limiting Ccl2, Cxcl1, and Cxcl10 expression by inducing A20 expression.

**Figure 4:**
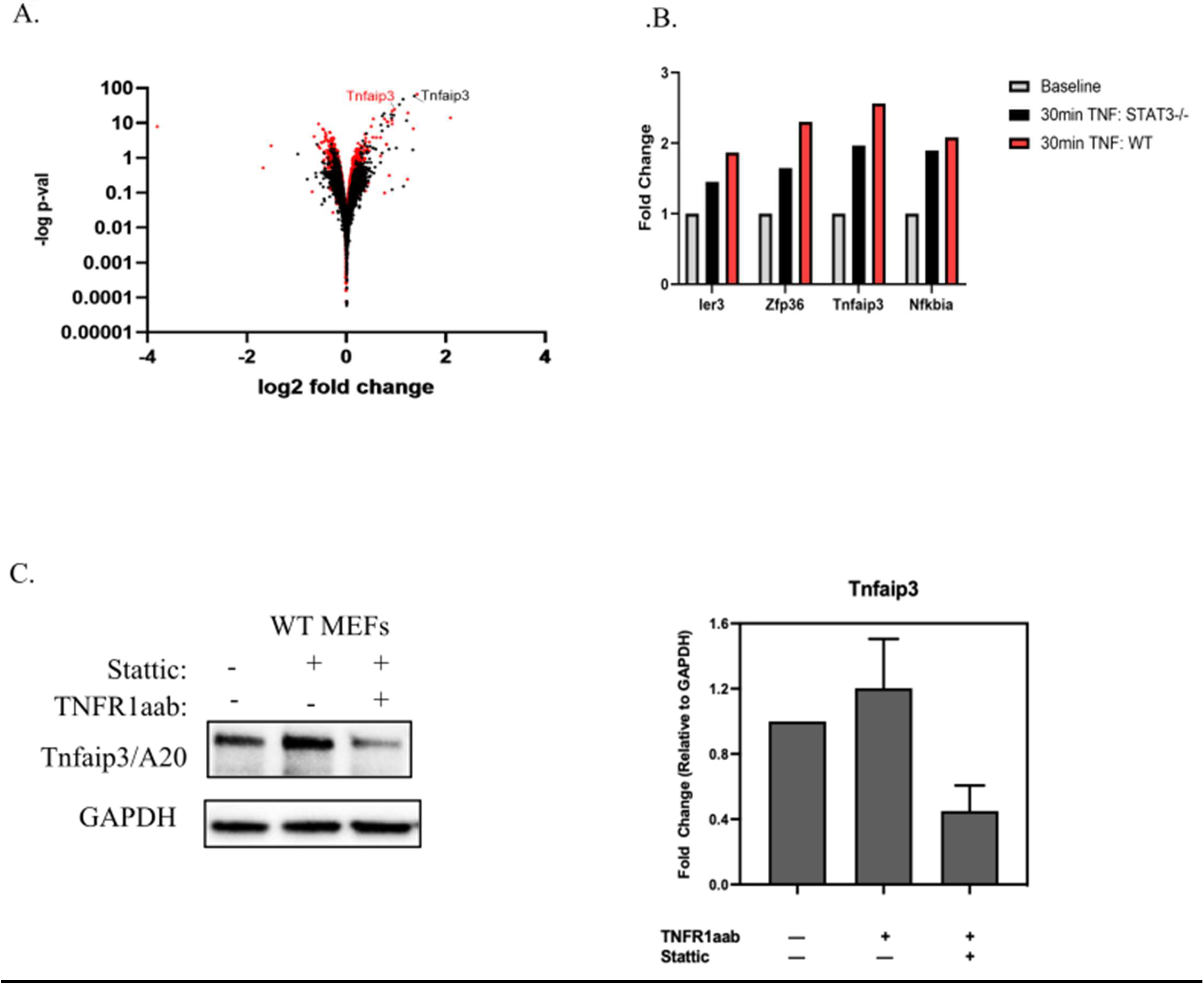
Negative TNFR1 regulator TNFAIP3/A20 is a STAT3 target gene A. Volcano plot showing the differentially expressed genes in WT and matched STAT3−/− MEFs stimulated with TNFα for 30 min as measured by RNA-sequencing. B. Genes in (A) that were induced to a greater extent in *Wt* than in *Stat3−/−* MEFs C. Western blot of Wt MEFs stimulated for 4 hours with TNFR1aab in combination with DMSO or Stattic (STAT3 inhibitor). Quantification of the band intensity is shown in the bar graph and normalized to GAPDH. The error bars represent the standard deviation of three independent biological replicates.

To test this hypothesis, exogenous A20/Tnfaip3 was expressed from a constitutive (rather TNF inducible) promoter by viral transduction of *Stat3−/−* MEFs. These cells were stimulated with TNFR1aab, and the secreted cytokines were assayed. A20 overexpression rescued the increased Ccl2, Cxcl1, and Cxcl10 expression observed in the parental STAT3−/− MEFs (Figure 5).

**Figure 5:**
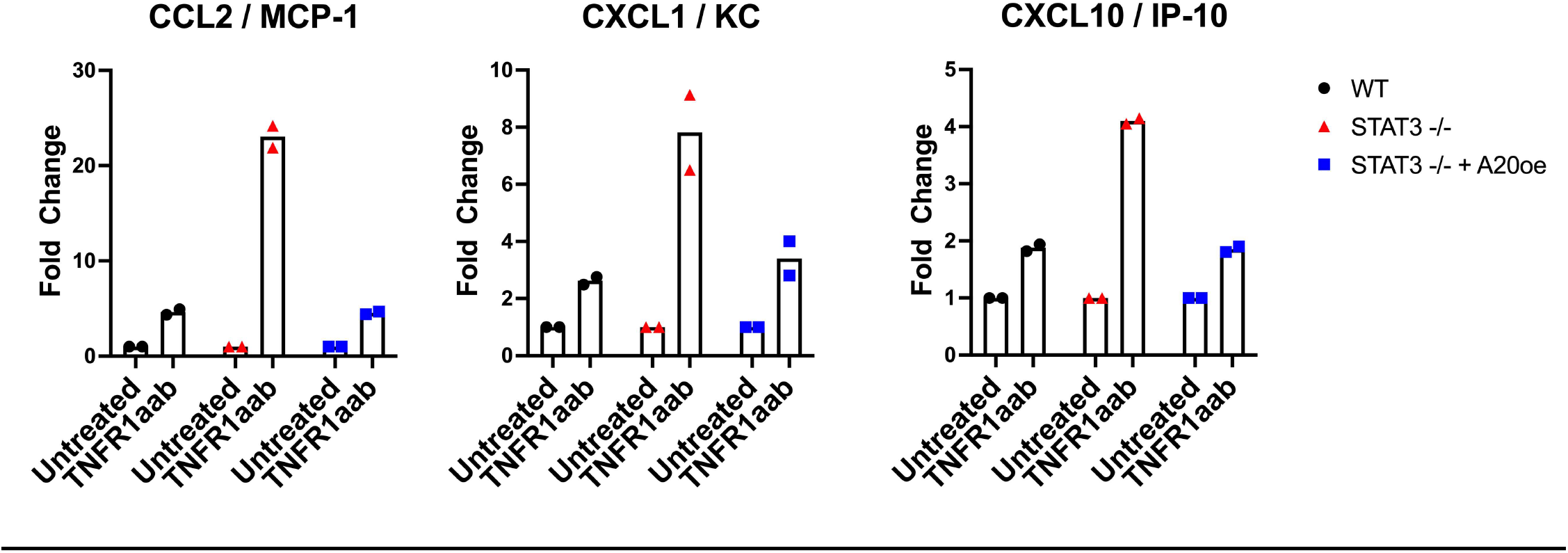
Effect of A20 expression on cytokine expression STAT3−/− MEFs. WT, STAT3−/−, or STAT3−/− MEFs transduced with a constitutive A20 expression plasmid were stimulated with the TNFR1aab antibody for 4 hrs and then subjected to the same cytokine panel described in Fig 3 and the Methods.

## Discussion

Inflammation plays a pivotal role in response to infection, injury, and trauma (1–3). The severity of the induced inflammatory response is important in many disease states and may determine whether the disease resolves or becomes chronic (4). TNFα has a pivotal role in the initiation and amplification of the inflammatory cascade; it is involved in regulating the release of chemokines and cytokines, oxidative stress, recruitment of immune cells and adhesion molecules, apoptosis, wound healing, and tissue-specific repair mechanisms. (*39*) Aberrant TNFα production and TNF receptor signaling have been associated with the pathogenesis of several diseases in which inflammation is an underlying element, including rheumatoid arthritis, Crohn’s disease, atherosclerosis, psoriasis, sepsis, diabetes, and obesity.(*40–43*) TNFα also orchestrates a cytokine cascade in many inflammatory diseases and because of its role as a “master-regulator” of inflammatory cytokine production it is a therapeutic target in such diseases.(*44*) Indeed, anti-TNFα drugs are licensed for treating several inflammatory diseases including rheumatoid arthritis and inflammatory bowel disease. (*45*)

Given that overactive TNFR1 signaling contributes to many diseases, there are important homeostatic mechanisms to ensure that TNFR1 signaling is transient and able to resolve the inflammatory response to prevent pathological inflammation. Indeed, several genes encode negative feedback regulators of the inflammatory process that are induced by the TNFR1 pathway, including NF-κB dependent expression of IkBα and A20.(*46, 47*) Knockout of these negative feedback regulators of TNF induced inflammation induces hyperinflammatory phenotypes in mice.

Since most of the genes downstream of TNFR1 were induced to a greater extent in the STAT3−/− than WT MEFs 4hrs post TNF stimulation, we hypothesized that STAT3 regulates the expression of a negative regulator of the TNFR1 pathway (Fig1). STAT3 can either promote or limit inflammation depending on the stimulus. For example, as noted earlier STAT3 is pro-inflammatory downstream of IL-6, but an effector of anti-inflammatory IL-10. (*27,28*) The work here shows that STAT3 plays an anti-inflammatory role downstream of TNFR1 in MEFS. Indeed, RNA sequencing 30 min after TNFR1 stimulation showed that STAT3 is necessary for full induction of A20 which limits the inflammatory response downstream of TNFR1 (see model Fig. 6).

**Figure 6:**
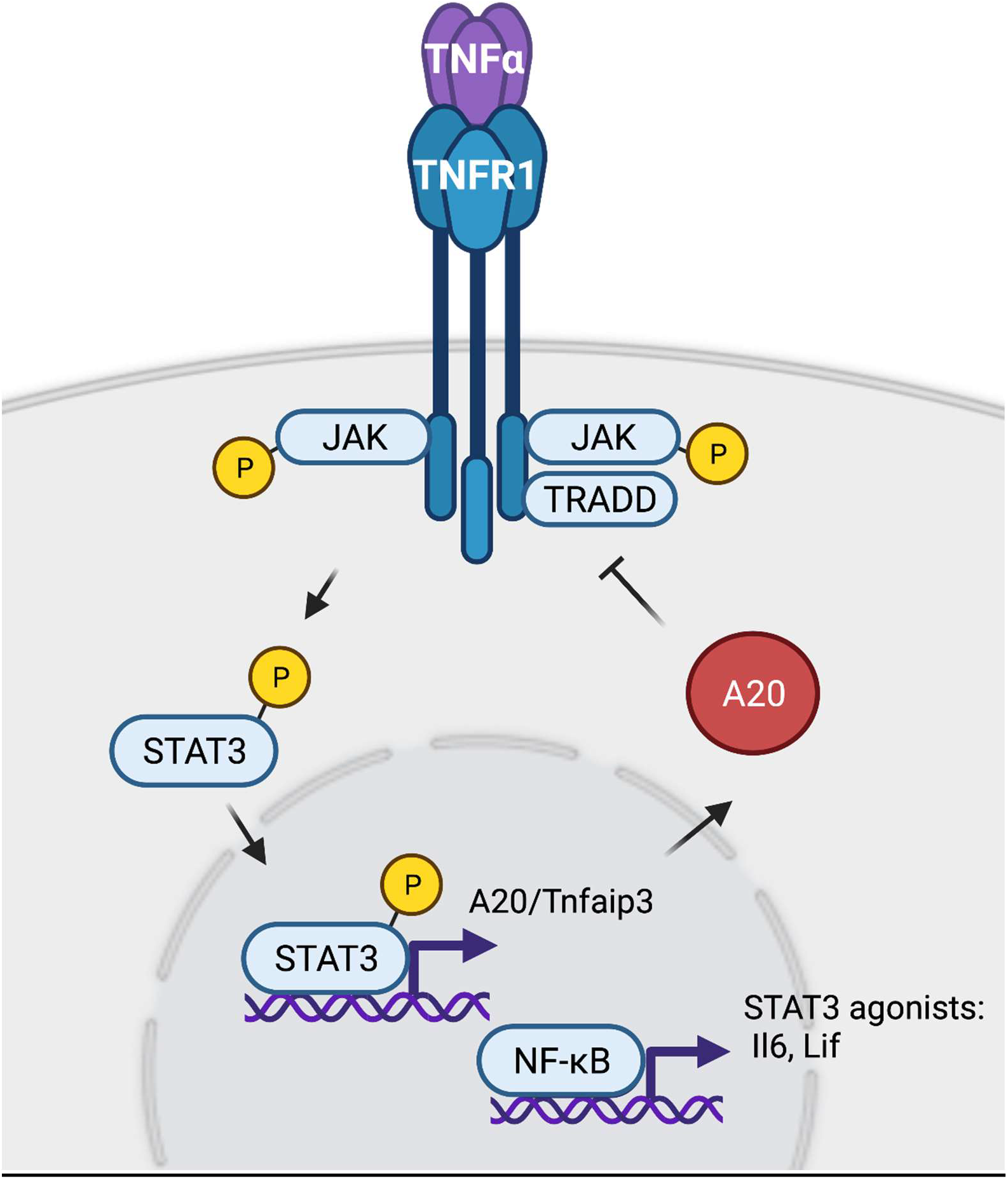
Model of the mechanism whereby STAT3 limits cytokine expression downstream of TNFR1. In previous work we showed that Jak2 (and cSRC) bind to TNFR1. Activation of TNFR1 brings receptor bound Jak2 proteins into proximity such that transphosphorylation and activation of Jak2 may take place. Activated Jak2 phosphorylates and activates STAT3 which enters the nucleus where it’s interaction with promoter binding sites results in the induction of A20 expression. A20 suppresses inflammatory cytokine expression through inhibition of NF-kB. The complexity of the inflammatory process is illustrated by the capacity of NF-kB to induce not only inflammatory cytokines but also STAT3 agonists such as IL6 and LIF. Image produced using *Created with BioRender.com*

Our observations identify a TNFR1-STAT3-A20 signaling pathway in MEFs. Most cell types express low levels of the A20 protein, which is induced by TNF in an NF-κB dependent manner. (46,47) A20 is deubiquitinating enzyme that is suppresses cytokine expression thus imposing a brake on pro-inflammatory processes mediated by TNF-α (37). Future studies are required to determine how STAT3 regulates A20. A20 is regulated by NF-κB (46,47) and STAT3 is co-operates with NF-κB in certain contexts at gene promoters (27,48). Therefore, an interesting future direction will determine whether and how STAT3 and NF-κB might co-operate at the A20 promoter to modulate its transcription.

Our observations are resonant of observations made with immune cells in which STAT3 suppresses signaling by Toll-like receptors, including TRL4, in phagocytes. (for review see 49). Stat3-deficient macrophages, neutrophils and dendritic cells produce elevated amounts of pro-inflammatory cytokines upon activation of TLR4. STAT3 inhibits the expression of the E2 ubiquitin-conjugating enzyme, Ubc13, required for TLR4 signaling and through this process impairs cytokine expression. Thus, a theme in inflammatory signaling may be its control by modulation of ubiquitination.

While most of the genes induced by TNFR1 were induced to a greater extent in the STAT3−/− than Wt MEFs, there were exceptions. This includes one of the cytokines validated at the protein level, GM-CSF. STAT3 also promotes GM-CSF expression in concanavalin A activated mesenchymal stromal cells (50). GM-CSF promotes the proliferation, differentiation, and activation of monocytes, granulocytes, macrophages, and dendritic cells *in vivo* and plays an important role in inflammatory cytokine responses. (51) Overall, the data presented here show that TNFR1 activates STAT3 which suppresses inflammatory cytokine production through A20 expression thereby diminishing cytokine expression and through GM-CSF expression which mobilizes the development of immune cells necessary for the host response to invasive stimuli.

## Materials and Methods

### Cell Culture

Wild type (Wt) and *Stat3−/−* mouse embryonic fibroblasts (MEFs), a kind gift from Dr. Albert Baldwin, were maintained in RPMI 1640 (Gibco) supplemented with 10% fetal bovine serum (Gibco) and 1X Penicillin-Streptomycin-Glutamine (Gibco). The WT MEFs used in the validation experiments were a kind gift from the National Institutes of Health (NIH, Zhenggan Liu at the Cell and Cancer Biology Branch, National Cancer Institute).(*58*) HEK293T cells, a gift from Dr. Hassan Alaoui (Department of Surgery, UCSF), were maintained in DMEM (Gibco) supplemented with 10% fetal bovine serum and 1X Penicillin-Streptomycin-Glutamine.

### RNA Sequencing

Total RNA was isolated using NucleoSpin mini RNA kit (Macherey-Nagel, Düren, Germany) according to the manufacturer’s protocol. RNA quality control, library preparation and Illumina sequencing were performed by Novogene Corporation Inc. (Sacramento, CA, USA). Raw sequencing data preprocessing, mapping to the reference genome (mm10) and gene expression quantification was performed by Novogene. Differential gene expression analysis was performed by using the iDep tool .(*59*) ENRICHR tool was used for pathway analysis(*32–34*)

### Plasmid constructs and viral transduction

Stable A20 overexpression in *Stat3−/−* MEFs was achieved by lentiviral transduction. Transfer plasmid containing the A20 insert was obtained from Origene (MR210582L4, Tnfaip3 NM_009397, OriGene Technologies, Inc.MD, USA). Packaging (psPAX2) and envelope plasmids (pMD2.G) were a gift from Dr. Hassan Alaoui. Transfer and packaging vectors were transfected into HEK 293T cells to produce lentiviruses using Lipofectamine 3000 reagent (ThermoFisher Scientific, MA, USA). Lentiviruses were harvested 48hr post transfection, 10X concentrated using Lenti-X concentrator (Takara Bio Inc, Japan), and used to infect *Stat3−/−* MEFs in the presence of Transdux Max (System Biosciences, Palo Alto, CA, USA) reagent. Fresh medium containing puromycin (Invitrogen, MA, USA) was added 24 hours later and the cells were maintained and selected for 2 weeks. A20 overexpression in *Stat3−/−* MEFs was confirmed by immunoblotting (data not shown).

### Multiplex Cytokine Assays

MEFs were grown in on 10 cm dishes until 80% confluent, serum-starved for 24hrs and stimulated with TNFR1aab (R&D Systems, MN, USA) for 4 hrs. Samples of the media were then collected and analyzed in duplicate for cytokine secretion by using a multiplex fluorescent bead assay (Eve Technologies, Calgary, Alberta, Canada).

### Western Blotting

Cells were lysed with RIPA buffer (ThermoFisher Scientific), and lysate protein concentrations were measured using Qubit protein assay kit (ThermoFisher Scientific). For Western blots, 30–50 μg of protein was resolved in 4-20% TGX gel (Bio-Rad Laboratories, Inc., Hercules, CA, USA) under reducing conditions and transferred onto a nitrocellulose membrane (Bio-Rad Laboratories). The following primary antibodies were added after blocking for an hour with 5% non-fat milk: STAT3 and Tyr705 pSTAT3 (Cell Signaling Technology, MA, USA), GAPDH (ThermoFisher Scientific), A20/TNFAIP3 (Cell Signaling Technology). Horseradish peroxidase-conjugated secondary antibody (Bio-Rad Laboratories) was then used, followed by visualization with FluoroChem digital imaging system (ProteinSimple Inc., CT, USA). Relative quantification of protein bands was performed using ImageJ software (NIH).

